# Perturbation of PTEN-PI3K/AKT Signalling Impaired Autophagy Modulation in Dystrophin-Deficient Myoblasts

**DOI:** 10.1101/125476

**Authors:** Muhammad Dain Yazid, Janet Smith

## Abstract

Alteration of single protein regulation has given a massive implication in Muscular Dystrophy pathogenesis. Herein, we investigated the contribution of defected dystrophin that has impaired PI3K/Akt signalling and subsequently reduced autophagy in dystrophin-deficient myoblasts. In this study, dfd13 (dystrophin-deficient) and C2C12 (non-dystrophic) myoblasts were cultured in low mitogen condition for 10 days to induce differentiation. Analyses of protein expression has been done by using immunoblot technique, immunofluorescence and flow cytometry. In our myoblasts differentiation system, the dfd13 myoblasts did not achieved terminal differentiation as fewer myotube formation and fast-myosin heavy chain expression almost not detected. Immunoblot analysis showed that PTEN expression is profoundly increased in dfd13 myoblasts throughout the differentiation day. As a result, the PI3K activity is decreased and has caused serine/threonine kinase Akt inactivation. Both residues; Thr308 and Ser473, on Akt were found not phosphorylated. The mTOR activation by Ser2448 phosphorylation was decreased indicates an impairment for raptor and rictor binding. Unable to form complexes; mTORC1 target protein, p70S6K1 activation was found reduced at the same time explained un-phosphorylated-Akt at Ser473 by rictor-mTORC2. As one of Akt downstream protein, transcription factor FoxO3 regulation was found impaired as it was highly expressed and highly mainly localised in the nucleus in dfd13 towards the end of the differentiation day. This occurrence has caused higher activation of autophagy related genes; Beclin1, Atg5, Atg7, in dfd13 myoblasts. Autophagosome formation was increased as LC3B-I/II showed accumulation upon differentiation. However, ratio of LC3B lipidation and autophagic flux were shown decreased which exhibited dystrophic features. As a conclusion, destabilisation of plasma membrane owing to dystrophin mutation has caused the alteration of plasma membrane protein regulation particularly PTEN-PI3K, thus impaired autophagy modulation that critical for myoblasts development.

## Introduction

The absence of dystrophin has a massive impact on myoblast structure. The destabilisation of the plasma membrane in dystrophin-deficient myoblasts from an *mdx* mouse (Duchenne muscular dystrophy (DMD) model) affects transmembrane protein stability, as well as the protein-anchored cytoplasmic layer of the cell membrane thus changing signalling. One of the conserved signalling pathways for skeletal muscle differentiation is the PI3K/Akt/mTOR pathway. The regulation is via insulin-like growth factor-1 receptor (IGF-1R) activation following IGF1 binding. IGF-1R plays a major role in the regulation of skeletal muscle growth and many researchers have reported on the impairment of the Igf-1 pathway and its downstream protein regulation in Muscular Dystrophy (MD).

Muscular dystrophy is inherited muscle diseases caused by a genetic disorder, i.e. deletion, duplication or a point mutation of the *dmd* gene (dystrophin) which is located on the human X chromosome (1). These diseases are characterised by progressive skeletal muscle weakness, defects in muscle proteins, and the death of muscle tissue (2). A person suffering from the disease will gradually lose the ability to perform daily routine activities, starting as early as after birth to 2 years old, and by around the ages of ten years old they will need a wheelchair for movement, with death occurring in their early 20s, mostly due to cardiomyopathy together with respiratory failure (3,4). The life expectancy for each individual who has MD depends on the degree of weakened muscles in the heart and lungs. However, there remains no treatment to cure MD, either drugs based or reverse genetic mutations. Currently, drug therapy, i.e. corticosteroids, is used to slow muscle degeneration, while physical therapy, speech therapy and orthopaedic appliances are used to support MD patients in their daily lives (5).

The PTEN-PI3K/Akt signalling pathway regulates multiple biological processes, such as cell proliferation, differentiation, autophagy and apoptosis. A disruption in this pathway can cause impairments to these processes, as can occurs in cancers due to an increase in Akt activity caused by the down-regulation of PTEN expression which is affected by PI3K activation. However, there is a different scenario in dystrophin-deficient myoblasts where PTEN expression is significantly higher and extensively expressed upon differentiation. PTEN, also known as a tumour suppressor, is mutated within cancer cells. PTEN acts as a lipid phosphatase and hydrolyses phosphates in the 3′ position of phosphoinositides. Previous studies have found that *PTEN* mRNA is elevated in the muscle of *mdx* mice (6,7). The elevation of PTEN expression and its activity due to the deregulation of the PI3K/Akt signalling pathway in dystrophin-deficient dog muscle has also been shown in the GRMD dog (8).

Many studies have been carried out concerning PI3K/Akt impairment in the *mdx* mouse as well as in DMD in humans. Activation of Akt has been found to be higher in the *mdx* mouse (6,9) and also in DMD patients (10). As Akt plays a major role in cell signalling, its impairment leads to various defects in downstream protein signalling, including cell proliferation, differentiation, apoptosis, autophagy and protein synthesis perturbation. Akt has also been reported to be highly activated and associated with α-integrin in the *mdx/utr^-/-^* mouse model (11). Boppart et al. (2011) demonstrated that α-integrin transgenic *mdx/utr^-/-^* mice (α7βX2-*mdx/utr-/-*) showed increased phosphorylation of Akt at Ser473, indicating that α-integrin expression is connected to Akt. The *mdx/utr^-/-^* mouse lacks both dystrophin and utrophin, and develops a severe pathology that closely resembles that seen in DMD (11).

Similarly, PTEN has been reported to contribute to the MD phenotype. A study by Feron et al. (2009) on the Golden Retriever muscular dystrophy (GRMD) dog demonstrated that PTEN was present at high levels, which led to a reduction of Akt1, glycogen synthase kinase-3β (GSK3β) and p70S6K, in addition, ERK1/2 displayed decreased phosphorylation levels in GRMD dog muscle. The GRMD dog is characterised by rapidly progressive clinical dysfunction, severe muscle weakness, and displays a disease progression that is more similar to human DMD compared to the *mdx* mouse (8).

Autophagy is the process of engulfment of cargo into the double lipid formation known as a autophagosome, which eventually fuses with a lysosome and is degraded (12). Autophagic activity is rapidly increased in cells under stress conditions and nutrient deprivation in order to maintain cell homeostasis (13). Generally, defective autophagy exhibits a dual response; at high levels, it causes muscle atrophy, whilst at low levels it contributes to muscle degeneration. High levels of autophagy result in excessive protein degradation due to high levels of autophagy gene activation (13). In this state inactivation of Akt inhibits the activation of autophagy related genes indirectly via mTOR and/or directly phosphorylates the FoxO3a transcription factor (14). This event leads to reduced muscle mass and muscle wasting. In contrast, low level autophagy is defined as low autophagy activity which causes an accumulation of dysfunction and unused organelles, i.e. mitochondria, as well as unfolded protein within skeletal muscle. This condition leads to an altered muscle structure with prominent myonuclei centralisation and fusion abnormalities, which eventually weakens the muscle leading to the dystrophic phenotype (15). Therefore, maintaining autophagy at the appropriate level is crucial within skeletal muscle.

Autophagy is defective in *mdx* mice (16) and DMD humans (17). Restoration of beclin1 levels in *Col6a1^-/-^* animals and long-term exposure to a low-protein diet (9,13) can reactivate autophagy and partly ameliorate the dystrophic features/phenotype. *Col6a1^-/-^* animals display an impairment of basal autophagy, which determines the persistence of dysfunctional organelles in muscle fibres leading to muscle degeneration (13). Treatment with a long-term low-protein diet can reactivate autophagy by normalised Akt activation, thus increasing LC3B conversion and up-regulating autophagy related genes (17).

Over a decade, many researchers have reported on impairment of Igf-1 pathway and its downstream protein regulation in muscular dystrophy. However, to further understand the impaired regulation of autophagy in dystrophin-deficient myoblast, the protein profiling of this pathway has been determined which sequentially provide information on autophagy modulation and activation. At the beginning, we reported the capacity of dystrophin-deficient myoblasts differentiation via analysis of myotube formation and the expression of specific myosin heavy chain as a terminal differentiation marker. Furthermore, we showed the elevation of PTEN perturbed PI3K/Akt/mTOR signalling thus excessively activate autophagy related genes via FoxO3 but subsequently showed reduction of autophagic flux in dystrophin-deficient myoblast. Overview of this event as illustrated in at the end of this article.

## Materials and methods

### Cell culture and differentiation

The C2C12 myoblast cell line was used in this study, and was established from an adult mouse myoblast C2 cell line derived from the thigh muscle of a 2-month-old mice. The dfd13 cell line was derived from a 5-week-old *mdx* mouse (18,19). Mouse embryonic fibroblast (MEF) cells were a gift from Adil Rashid, University of Birmingham, and were used as a control for the phosphorylation of Akt at threonine-308. Both myoblasts were maintained in growth medium (GM) consist of Dulbecco’s Modified Eagles Medium (DMEM) (Invitrogen Ltd, UK) supplemented with 10% Fetal Bovine Serum (FBS) (Sigma-Aldrich, UK), 1% penicillin/streptomycin (Sigma-Aldrich, UK) and 1% L-Glutamine (Sigma-Aldrich, UK) before plated for differentiation. To differentiate myoblasts, GM was replaced with differentiation medium (DM) which is DMEM supplemented with 2% Horse Serum (Sigma-Aldrich, UK), 1% penicillin/streptomycin (Sigma-Aldrich, UK) and 1% L-Glutamine and cultured for 10 days.

### Total protein extraction

Protein was extracted from cells cultured in a 10 cm dish. The medium was removed and the cells washed twice with PBS before adding approximately 300 μL of lysis buffer (0.5% triton X-100, 0.5% deoxycholic acid; 0.5 M NaCL; 0.02 M Tris; 0.01 M EDTA, pH 7.5) containing a protease inhibitor (complete ULTRA tablets, EDTA-free, protease inhibitor cocktail, Roche, UK). The cells were scrapped and collected into a tube before being subjected to centrifugation at 14000 rpm for 15 minutes at 4 °C. The pellet was discarded and the supernatant transferred to a new tube and stored at −20 °C.

### Protein sub-fractionation

The rapid efficient and practical (REAP) protocol was used for protein sub-fractionation (20). All reagents were chilled and kept on ice all times. The medium was removed and the cells washed three times with PBS before adding approximately 1 mL of ice-cold PBS. The cells were scrapped and collected into a new tube and pop-spun for 10 seconds and the supernatant discarded. The pellet was resuspended in 900 μL of ice-cold PBS containing 0.1% nonyl-phenoxypolyethoxylethanol (NP-40) and approximately 300 μL was removed to a new tube labelled as whole cell lysate (WCL). The remaining cells were pop-spun for 10 seconds and approximately 300 μL was transferred to a new tube labelled as the cytosolic fraction, while the remainder of the supernatant was discarded. The pellet was washed with 1 mL of PBS containing 0.1% NP-40) and pop-spun for 10 seconds; this step was repeated 2 to 5 times to eliminate the cytosolic fraction. After washing was completed this was considered to be the nuclear fraction. The cytosolic fraction was also pop-spun 3 times to separate the remaining nuclear fraction. Both the WCL and nuclear fraction were sonicated three times for 10 seconds and boiled for 1 minute.

### Western blotting

Equal protein loading was prepared before resuspend with 2 × Laemmli sample buffer. The mixture was collected by short spin centrifugation and boiled for 5 minutes before loaded into SDS-polyacrylamide gel electrophoresis. Lysate were separated on SDS-polyacrylamide gel electrophoresis (6, 8 or 15% acrylamide) with 200 V for 1 hour. The separated protein were transferred onto nitrocellulose membrane using Transblot Turbo^®^ Transfer System (BioRad, UK). The membranes were blocked in 5% reduced fat milk for 1 hour prior to incubate with primary antibodies. The antibodies are Fast-MyHC, α-tubulin, SSRP-1 (1:1000) and Desmin, (1:2000), from Sigma-Aldrich, UK. PTEN (1:1000) (a gift from Dr. Zubair Ahmed, Medical School, University of Birmingham). PI3K, phospho-PI3K (Tyr458), Akt, phospho-Akt (Ser473), mTOR, phospho-mTOR (Ser2448), p70S6 Kinase, phospho-p70S6 Kinase, Rictor, phospho-rictor (Thr1135) and FoxO3a (1:1000) from Cell Signalling Technology, UK. Beclin1, Atg5, Atg7 and LC3B (1:1000) (a gift from Dr. Melissa Grant, School of Dentistry, University of Birmingham).

### Immunoflourescence

The cover slips were acid-etched for 5 minutes with nitric acid. This will help myoblasts firmly attached onto the cover slips prior for immunofluorescence. It has been sent for sterilisation before myoblasts were seeded onto it. The myoblasts/myotubes were fixed using 4% PFA. After fixation cells were permeabilised with 0.25% triton X-100 for 5 minutes to allow the antibody to bind specifically. The cells were then washed three times for 5 minutes with PBS before blocking with 5% BSA at room temperature for 30 minutes. The blocking solution was removed and the cells incubated with the primary antibody Fast-MyHC at 4°C overnight. The primary antibody was removed the next day and the cells washed before probed using a secondary antibody (biotinylated) for 1 hour at room temperature. After washing, the cells were incubated with streptavidin-Texas red (1:1000) at room temperature for 1 hour in the dark. The cells were then counterstained with DAPI (1:100) and mounted onto slides using DakoCytomation. The slides were wrapped in aluminium foil and stored at 4°C.

### Autophagy assay: cells labelling

An autophagy assay was performed using a CYTO-ID Autophagy Detection Kit (Enzo Life Science, Switzerland) and 1 × 10^5^/mL myoblasts were added to every well in a 24-well plate and maintained via standard tissue culture practice. Cells were grown in a humidified incubator at 37 °C and 5% CO_2_; cells were not allowed to become over-crowded and the density should not exceed 1 × 10^6^ /mL. In this experiment 60 mM of chloroquine was used as positive control, while unstained cells were utilised as a negative control. Myoblasts were treated with chloroquine for 4 hours in a humidified incubator at 37 °C and 5% CO2. At the end of the treatment cells were trypsinised and collected into a fresh 1.5 mL microtube. Cells were subjected to centrifugation for 5 minutes at1000 rpm at room temperature and washed by re-suspending in 1x assay buffer and the centrifugation step repeated. Each cell pellet was then re-suspended in 250 μL of 1x assay buffer. Approximately 250 μL of the diluted CYTO-ID Green stain solution was added to all the cells except the negative control, and incubated for 30 minutes at room temperature in the dark. The cells were collected by centrifugation, washed with 1x assay buffer, and the pelleted cells re-suspended in 500 μL of 1x assay buffer. The samples were analysed using the green (FL1) channel of a flow cytometer.

### Flow cytometer

Experiments were performed using a FACS Analyser CyAn B (Beckman Coulter, USA) at the Institute of Biomedical Research, University of Birmingham. Samples were run using Summit V4.3 software and all data were saved in the fcs format developed by the Society of Analytical Cytology, thereby allowing further analysis using other software packages. Data were analysed using the software FCS Express 6 Plus (De Novo Software, USA) for three parameters; side scatter (SS), forward scatter (FS) and Fluorescein Isothiocyanate (FITC). Gating was performed based on the size and complexity of the myoblasts as illustrated in density plots. Gated data were further analysed to count myoblasts labelled with FITC and the results presented in a histogram format.

### Statistical analysis

All Western blots and flow cytometry were repeated at least three times for each experiment. Statistical analyses were carried out using the student t-test (Microsoft Excel) and differences were considered significant at p<0.05 and p<0.01.

## Results

### Dystrophin-deficient myoblasts do not achieve terminal differentiation

*In vitro* differentiation of myoblasts can be induced in culture through the use of a low mitogen medium (2% horse serum) for a few days. In this study, both C2C12 (non-dystrophic) and dfd13 (dystrophin-deficient) myoblasts were cultured in low mitogen medium (differentiation medium; DM) for 10 days prior to determining their terminal differentiation capacity. Immunofluorescence analyses (Fig 1A-C) showed both non-differentiated and differentiated myoblasts. Multinucleated myotube formation can be clearly seen on day 10 of differentiation in C2C12 (Fig 1A) myoblasts but is hardly/rarely found in dfd13 myoblasts where less multinucleated cells were observed (Fig 1B). This indicates that dfd13 differentiation capacity is impaired; however, the differentiating dfd13 myoblasts can be seen to be aligned and seem to be ready for cell fusion to become myotubes.

**Fig 1.**
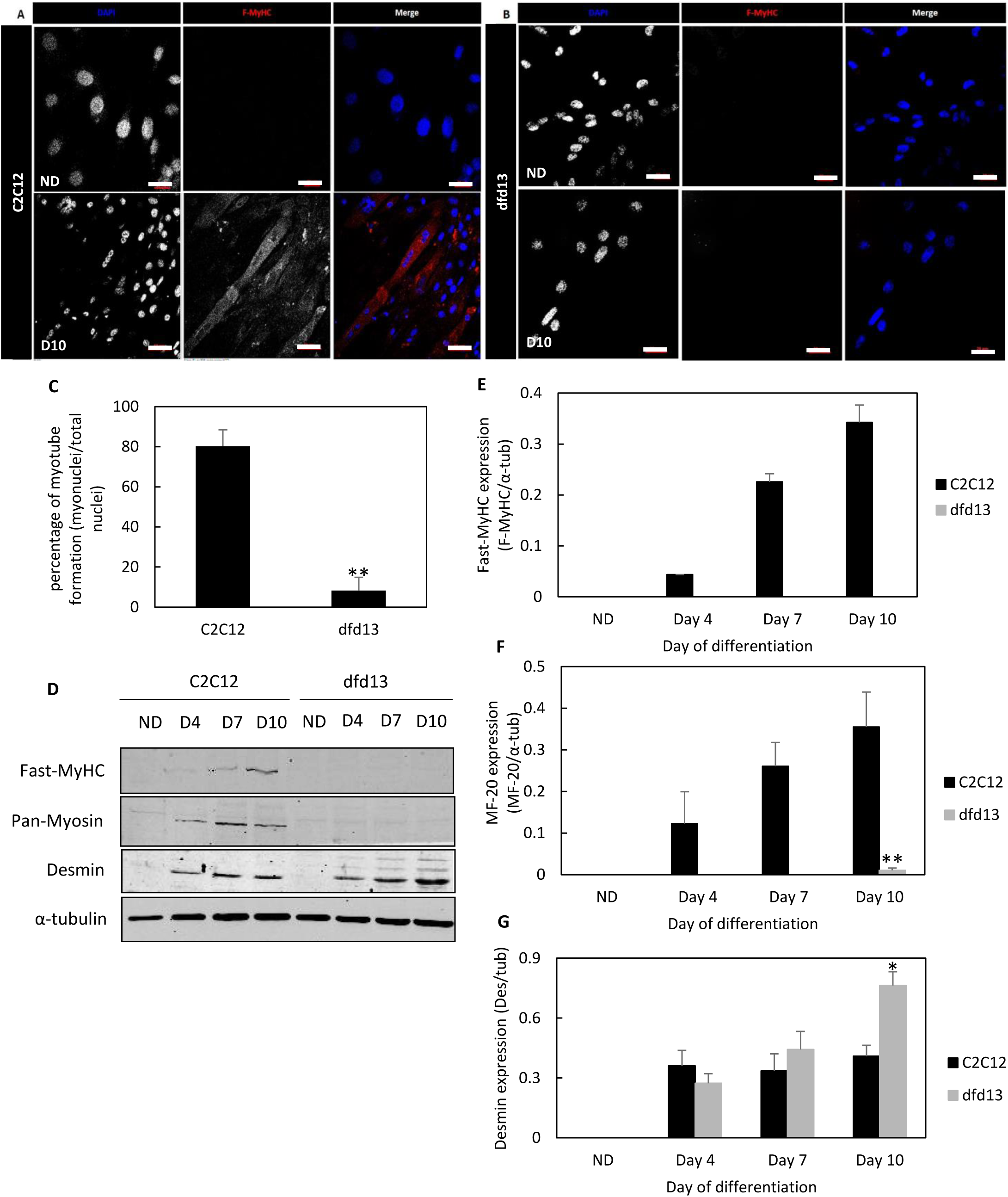
Dystrophin-deficient myoblasts have impaired differentiation capacity. Approximately 1.5 × 10^4^ of both C2C12 (non-dystrophic) and dfd13 (dystrophin-deficient) myoblasts were cultured in GM before being transferred to DM and allowed to differentiate for 10 days in a 48-well plate (on acid-etched cover slip). On day 10 cells were fixed in 4% PFA and stained with an anti F-MyHC antibody, while DAPI was used as a nuclear counterstain. Immunofluorescence analysis of F-MyHC in (A) C2C12 and (B) dfd13 cells during the non-differentiated stage and after 10 days of differentiation. (C) Percentage myotube formation was calculated by counting the nuclei present in myotubes (myonuclei) per total nuclei from 10 random microscope fields. Both C2C12 and dfd13 myoblasts were seeded in 10-cm dishes and cultured in GM until 80-90% confluent. The myoblasts were washed twice with PBS and then cultured in DM for 10 days, which was changed every 2 days. Total protein was extracted at the indicated time points and immunoblotted with antibodies recognising the proteins F-MyHC, pan-myosin (MF-20) and desmin. (D) Representative immunoblot of the proteins during myoblast differentiation with α-tubulin expression as a loading control. Densitometry analyses of (E) F-MyHC (MY-32), (F) pan-myosin (MF20) and (G) desmin expression. The graphs represent an average of three repeats from different samples. ND: non-differentiated; D10: day 10 of differentiation; **: significantly different (p<0.01) compared to C2C12 myoblasts. *: significantly different (p<0.05) compared to C2C12 myoblasts GM: growth medium (DMEM + 10% FCS); DM: differentiation medium (DMEM + 2% horse serum).

Myotube formation was determined by counting the myonuclei present in a myotube that expressed MyHC (fast-type II, F-MyHC) and then normalised against total nuclei. F-MyHC, also called ‘fast-twitch’ fibres, were chosen as the terminal differentiation marker based on the existence of a spectrum of fibre types, with type II being the most developed form of myosin: type 1 ↔ 1/2A ↔ 2A ↔ 2A/2X ↔ 2X ↔ 2X/2B ↔ 2B (21). The number of myonuclei in C2C12 myotubes was 9-fold higher compared to myonuclei in differentiating dfd13 myoblasts (p<0.01; p = 2.9×10^−3^). The percentage of myonuclei per total nuclei was 80.2% ± 8.2 in C2C12 myotubes and 8.2% ± 6.6 in differentiating dfd13 myoblasts (Fig 1C).

Immunoblotting for F-MyHC, pan-myosin and desmin was performed on days 4, 7 and 10 of differentiation (Fig 1D). Generally, F-MyHC expression was increased upon differentiation in C2C12 myoblasts but none of the differentiating dfd13 myoblasts showed any expression (Fig 1E). However, pan-myosin was expressed at day 10 in dfd13 myoblasts (Fig 1F), and there was a significant difference in expression (p<0.01; p = 7.78 × 10^−3^) when compared to C2C12 myoblasts. Desmin expression in both myoblasts was also examined and was found to be drastically increased upon differentiation, in C2C12 myoblasts. In differentiating dfd13 myoblasts, desmin expression gradually increased over the 10 days and was significantly higher (p<0.01; p = 2.24 × 10^−3^) in comparison to levels in C2C12 myotubes (day 10) (Fig 1G). Desmin was used as an intermediate differentiation marker and is a muscle specific type II intermediate filament that integrates the sarcolemma, Z-disk and nuclear membrane in myoblasts. An *in vivo* study by Smythe et al. (2001) indicated that desmin expression is up-regulated during myogenesis and demonstrated that desmin(-/-) showed delays in myotube regeneration 5 days after transplantation when compared to the control where desmin was detected (22). Desmin also has been reported overexpressed at molecular level in DMD patient (23).

### PTEN-PI3K regulation is perturbed in differentiating dystrophin-deficient myoblasts

Generally, PTEN expression was increased upon differentiation in both types of myoblasts. However, the accumulation of PTEN was found to be higher in differentiating dfd13 myoblasts compared to C2C12 myoblasts (Fig 2A). The accumulation of PTEN throughout differentiation was not significant in C2C12 myoblasts but there were significant increases on day 7 (p<0.05; p = 4.30×10^−2^) and day 10 (p<0.05; p = 4.88×10^−2^) when compared to non-differentiated cells. Densitometry analysis showed that PTEN expression was significantly higher (p<0.05; p = 2.07×10^−2^) in dfd13 myoblasts compared to C2C12 in non-differentiated stage. The differentiating dfd13 myoblasts were also showed significantly higher compared to the differentiating C2C12 myoblasts (p<0.01; p = 4.81×10^−3^) on day 4, (p<0.05), day 7 (p = 1.29×10^−2^) and day 10 (p = 1.95×10^−2^) (Fig 2B).

**Fig 2.**
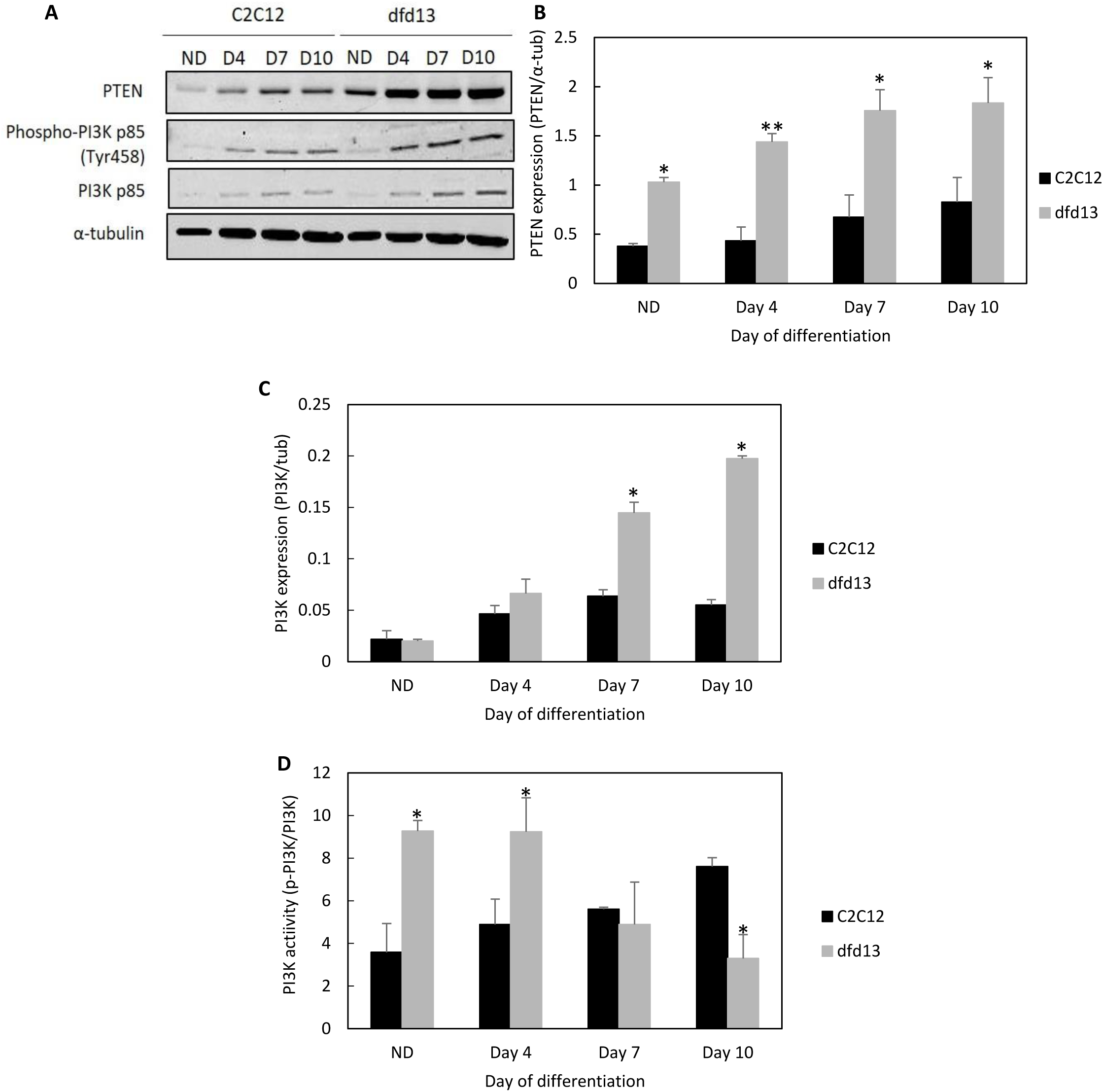
PTEN expression is higher while PI3K activation is decreased in dystrophin-deficient myoblasts. Myoblasts were cultured in GM until 80 to 90% confluent before washing twice with PBS and culturing in DM for 10 days with the media being changed every 2 days. Total protein extraction was performed at the indicated time points prior to immunoblot analysis. (A) Immunoblot analysis of PTEN, phospho-PI3K (p85) and total PI3K during myoblasts differentiation with α-tubulin expression as a loading control. Densitometry analysis of (B) PTEN expression, (C) total PI3K expression, and (D) PI3K activity. The graphs represent an average of three repeats from different samples. ND: non-differentiated; Significant different: * (p<0.05) and ** (p<0.01) compared to C2C12 myoblasts. GM: growth medium (DMEM + 10% FCS); DM: differentiation medium (DMEM + 2% horse serum)

PTEN has a negative effect on PI3K, as it concomitantly affects PI3K regulation. PI3K expression and its activity in both types of myoblasts was therefore examined. Generally, total PI3K expression was increased in dfd13 myoblasts throughout the differentiation period. However, significant accumulation of PI3K could be seen by day 7 (p<0.01; p = 1.04×10^−3^) and day 10 (p<0.01; p = 1.65×10^−3^) when compared to the non-differentiated dfd13 myoblasts. There was no significant difference in total PI3K expression in C2C12 myoblasts throughout the differentiation period compared to non-differentiated myoblasts. Densitometry analysis showed that there were significant differences in total PI3K expression in dfd13 myoblasts on day 4 (p<0.05; p = 1.70×10^−2^), day 7 (p<0.01; p = 2.08×10^−3^), and day 10 (p = 1.42×10^−3^) when compared to differentiating C2C12 myoblasts (Fig 2C). As illustrated in Fig 2D, densitometry analysis on PI3K activity was higher in dfd13 myoblasts at the non-differentiated stage (p<0.01; p = 9.89×10^−3^), day 4 (p<0.01; p = 1.69×10^−3^) and reduced at day 10 of differentiation (p<0.05; p = 3.25×10^−2^) when compared to C2C12 myoblasts.

In this study, phosphorylation of PI3K-p85 (regulatory subunit) was examined in order to observe PI3K activation. When cells are stimulated by the receptor, intrinsic tyrosine kinase phosphorylates p85 at Tyr458, causing a conformational change to p110 (the catalytic subunit) which increases its enzymatic activity for PIP2. From this data, it seems that high PTEN expression decreased PI3K activity, thus affecting its downstream protein activation.

### Akt is less activated in dystrophin-deficient myoblasts

As PTEN has been shown to be highly expressed in dfd13 myoblasts, it is predicted that Akt is not/less activated, as PTEN has previously been reported to modulate Akt activation in rhabdomyosarcomas cells; skeletal muscle cancer (24). In this study phosphorylation of Akt was not detected at Ser473 or Thr308 dfd13 myoblasts during differentiation (Fig 3A). It has been reported that phosphorylation of Akt by PDK1 at Thr308 partially activates Akt while full activation requires phosphorylation of Ser473 which can be catalysed by multiple proteins including rictor-mTORC2 (25). These activation is responsible for myoblast proliferation as well as differentiation.

**Fig 3.**
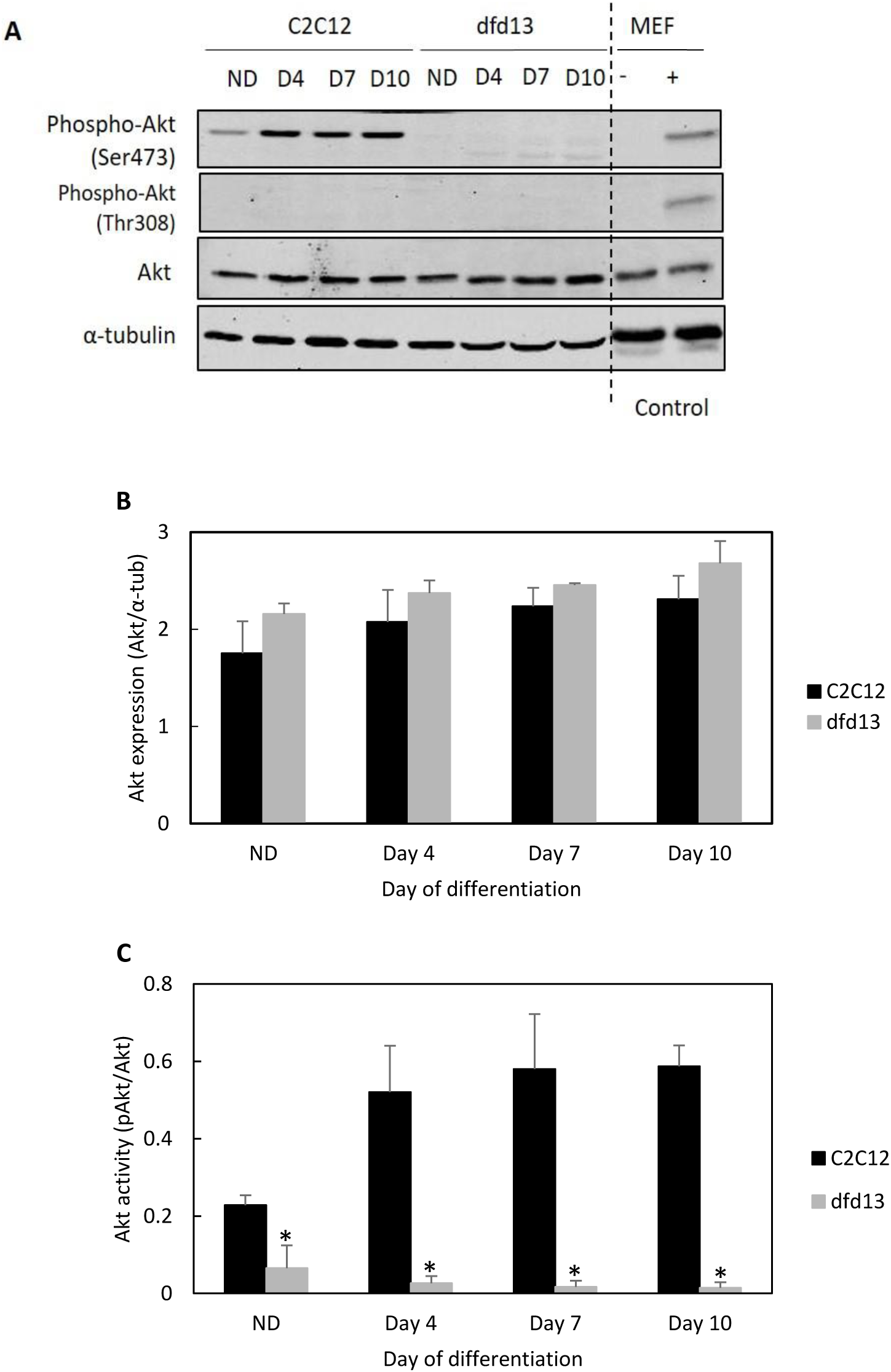
Akt is not/less activated in dystrophin-deficient myoblasts. Myoblasts were cultured in GM until 80 to 90% confluent before washing twice with PBS and culturing in DM for 10 days with the media being changed every 2 days. Untreated- and PDGF-treated MEF cells were used as a control for Akt phosphorylation at both Ser473 and Thr308. Total protein was extracted at the indicated time points prior to immunoblotting with antibodies recognising Akt, phospho-Akt (Ser473) and phospho-Akt (Thr308). (A) Immunoblot analysis during myoblast differentiation with α-tubulin expression as a loading control. Densitometry analysis of (B) Akt expression and (C) Akt activation at Ser473. The graphs represent an average of three repeats from different samples. ND: non-differentiated; +: PDGF-treated; -: non-treated; *: significantly different (p<0.05) compared to C2C12 myoblasts. GM: growth medium (DMEM + 10% FCS); DM: differentiation medium (DMEM + 2% horse serum).

As depicted in Fig 3B, total Akt expression increased in both types of myoblast during differentiation, with slightly higher levels present in dfd13 myoblasts (no significant different). Densitometry analysis of Akt activation via phosphorylation at Ser473 (Fig 3C) showed a significant difference when compared to C2C12 myoblast at all stages (non-differentiated p<0.05; p = 3.90×10^−2^, day 4 p<0.05; p = 1.17×10^−2^, day 7 p<0.05; p = 1.25×10^−2^ and day 10 p<0.01; p = 1.39×10^−3^). There was also a significant difference in Akt activation at Ser473 in C2C12 myoblasts when compared to the non-differentiated stage (day 4 p<0.05; p = 2.06×10^−2^ day 7 p<0.05; p = 3.31×10^−2^ and day 10 p<0.01; p = 1.41×10^−3^).

Akt is a downstream protein of PI3K. Its plays an important role and is recognised as one of the most critical pathways in the regulation of cell viability and maintenance in skeletal muscle mass (26). Therefore, its expression pattern in dystrophin-deficient myoblasts was investigated, as previously dfd13 myoblasts have been reported to undergo apoptosis when cultured in DM (27). As expected, the immunoblot data showed that Akt was not found/less activated in differentiating dfd13 myoblasts. Surprisingly, it was found that Akt was only activated by phosphorylation at Ser473 in C2C12 myoblasts. Akt was not phosphorylated at Thr308 in either type of myoblasts and consequently mouse embryonic fibroblast (MEF) cells were used as a positive control (gift from Adil Rashid, University of Birmingham) for this activation site.

### Rictor-mTORC2 is less activated in dystrophin-deficient myoblasts

A previous study reported that mTORC2 is essential for terminal myogenic differentiation (28). It was also reported that it participates in actin cytoskeleton arrangements, which might be involved in dystrophin functionality in myoblasts. Therefore, it was hypothesised that Akt inactivation is affected by rictor, a subunit of mTORC2, upstream of Akt and responsible for Ser473 phosphorylation. However, the upstream protein that regulates rictor-mTORC2 remains unclear. In this study both total rictor and mTOR expression was examined as well as its activation. However, it was impossible to classify the specific complexes of mTOR activity as phosphorylation at Ser2448 can cause the binding to both raptor and rictor (29).

As depicted in Fig 4A, immunoblot analysis of total rictor, mTOR and their phosphorylated forms was performed. Phosphorylated-rictor at Thr1135 was virtually not detected in dfd13 myoblasts throughout the differentiation period. However, rictor expression was increased upon differentiation in both types of myoblasts. Densitometry analysis showed that there was no significance difference in its accumulation in the two types of myoblasts (Fig 4B). Rictor expression only showed a significant increase (p<0.05; p = 2.75×10^−2^) in dfd13 myoblasts at day 10 compared to non-differentiated dfd13 myoblasts. Densitometry analysis of rictor activation by phosphorylation at Thr1135 (Fig 4C) showed a significant difference in C2C12 myoblasts compared to dfd13 myoblasts at all stages; non-differentiated p<0.05; p = 2.22×10^−2^, day 4 p<0.05; p = 3.52×10^−2^, day 7 p<0.05; p = 1.01×10^−2^ and day 10 p<0.01; p = 3.23×10^−3^).

**Fig 4.**
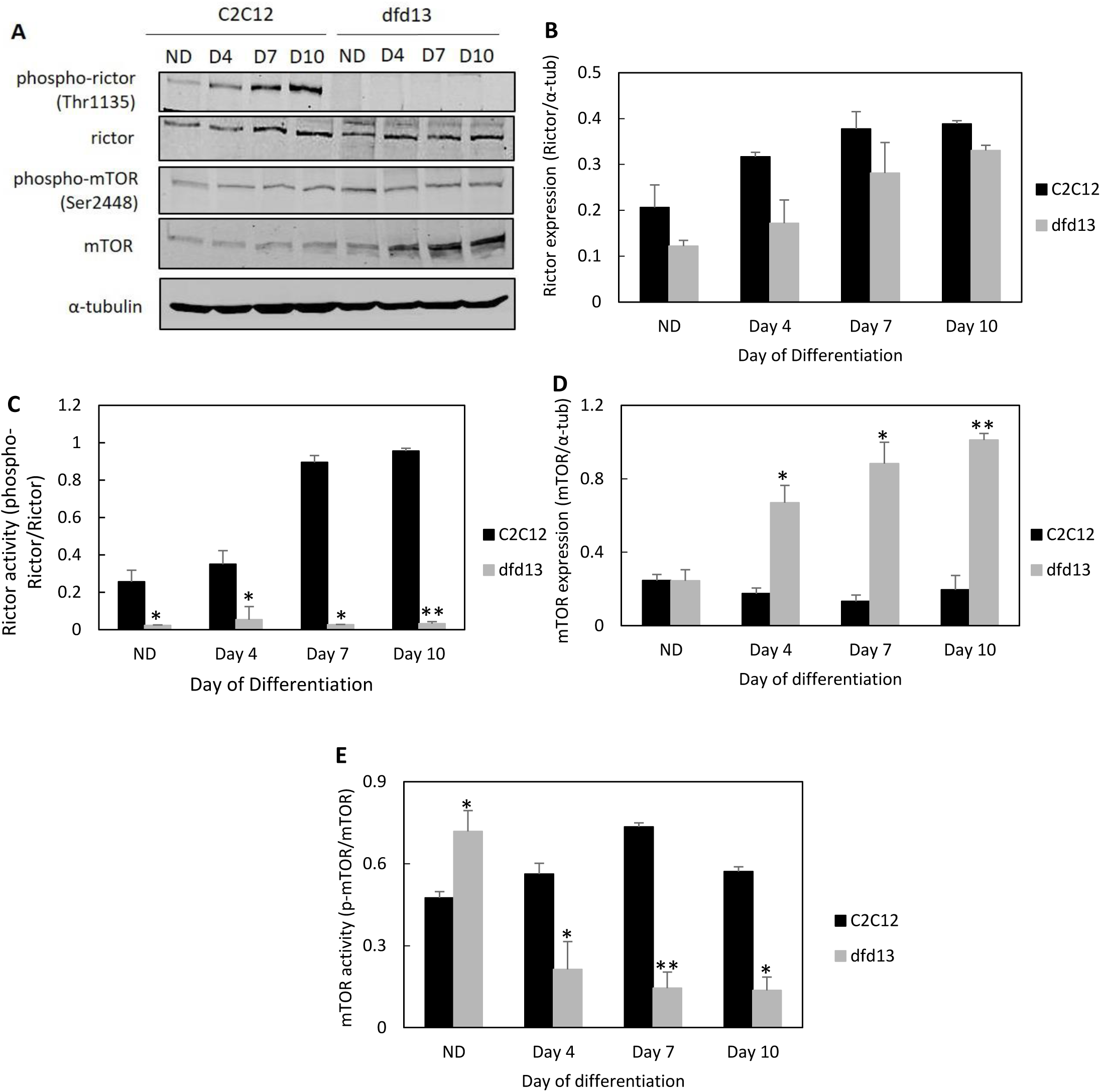
Rictor-mTORC2 activation is lower in dystrophin-deficient myoblasts. Myoblasts were cultured in GM until 80 to 90% confluent before washing twice with PBS and culturing in DM for 10 days with the media being changed every 2 days. Total protein was extracted at the indicated time points prior to immunoblotting with antibodies recognising phospho-rictor (Thr1135), rictor, phospho-mTOR (Ser2448) and mTOR proteins. (A) Immunoblot analysis during myoblast differentiation with α-tubulin expression as a loading control. Densitometry analyses of (B) total rictor expression (C) rictor activity (D) total mTOR expression, and (E) mTOR activity. The graphs represent an average of three repeats from different samples. ND: non-differentiated; *: significantly different (p<0.05) compared to dfd13 myoblasts. GM: growth medium (DMEM + 10% FCS); DM: differentiation medium (DMEM + 2% horse serum).

Fig 4D shows that total mTOR expression was increased in dfd13 myoblasts during differentiation but remained the same in C2C12 myoblasts. A significant accumulation was seen in dfd13 myoblasts on day 4 (p<0.05; p = 2.93×10^−2^), day 7 (p<0.05; p = 1.23×10^−2^), and day 10 (p<0.01; p = 8.55×10^−3^) when compared to C2C12 myoblasts. mTOR activity was significantly higher in non-differentiated dfd13 myoblasts when compared to non-differentiated C2C12 myoblasts (p<0.05; p = 4.15×10^−2^). Activity showed a significant reduction upon differentiation in dfd13 myoblasts while C2C12 myoblasts showed a significant accumulation when compared to dfd13 myoblasts on day4 (p<0.05; p = 2.23×10^−2^), day 7 (p<0.01; p = 6.78×10^−3^) and day 10 (p<0.05; p = 1.29×10^−2^) (Fig 4E).

From these results, it can be suggested that inactivation of Akt is caused by the inactivation of rictor in dfd13 myoblasts. This impairment is thought to effect other kinases, such as protein kinase C (PKC) and focal adhesion proteins, such as integrin-linked-kinase (ILK). Alterations to these kinases have been reported as one of the factors that causes MD. Expression and activation of mTOR was considered to represent endogenous levels of mTOR within both complexes i.e. mTORC1 or mTORC2, where distinct activation occurred. Phosphorylation of mTOR at Ser2448 makes it a major target for p70S6 kinase activation and is also an important event for raptor and rictor binding (29).

### Foxo3 expression is highly increased and predominantly localised in the nucleus of differentiating dystrophin-deficient myoblasts

Inactivation of Akt had an effect on another downstream protein, FoxO3 (Forkhead box O3), which is a target protein of Akt. FoxO3 is a transcription factor responsible for the activation of autophagy genes involved in the autophagy machinery. A previous study showed that FoxO3 controls autophagy in skeletal muscle *in vivo* and induced multiple autophagy genes, including LC3B transcription in skeletal muscle (30). In this study, FoxO3 expression in myoblasts was examined.

Immunoblot analyses (Fig 5A) showed that FoxO3 expression was increased in both types of myoblasts during differentiation. However, there was only a slight increase in FoxO3 in C2C12 myoblasts on day 4, and it remained at the same level until day 10. In non-differentiated dfd13 myoblasts FoxO3 levels were found to be significantly lower (p<0.01; p = 6.63×10^−3^) compared to C2C12 myoblasts. FoxO3 then increased throughout the differentiation period and there was a significant difference in dfd13 myoblasts (p<0.01; p = 4.45×10^−3^) on day 10 compared to C2C12 myoblasts (Fig 5B).

**Fig 5.**
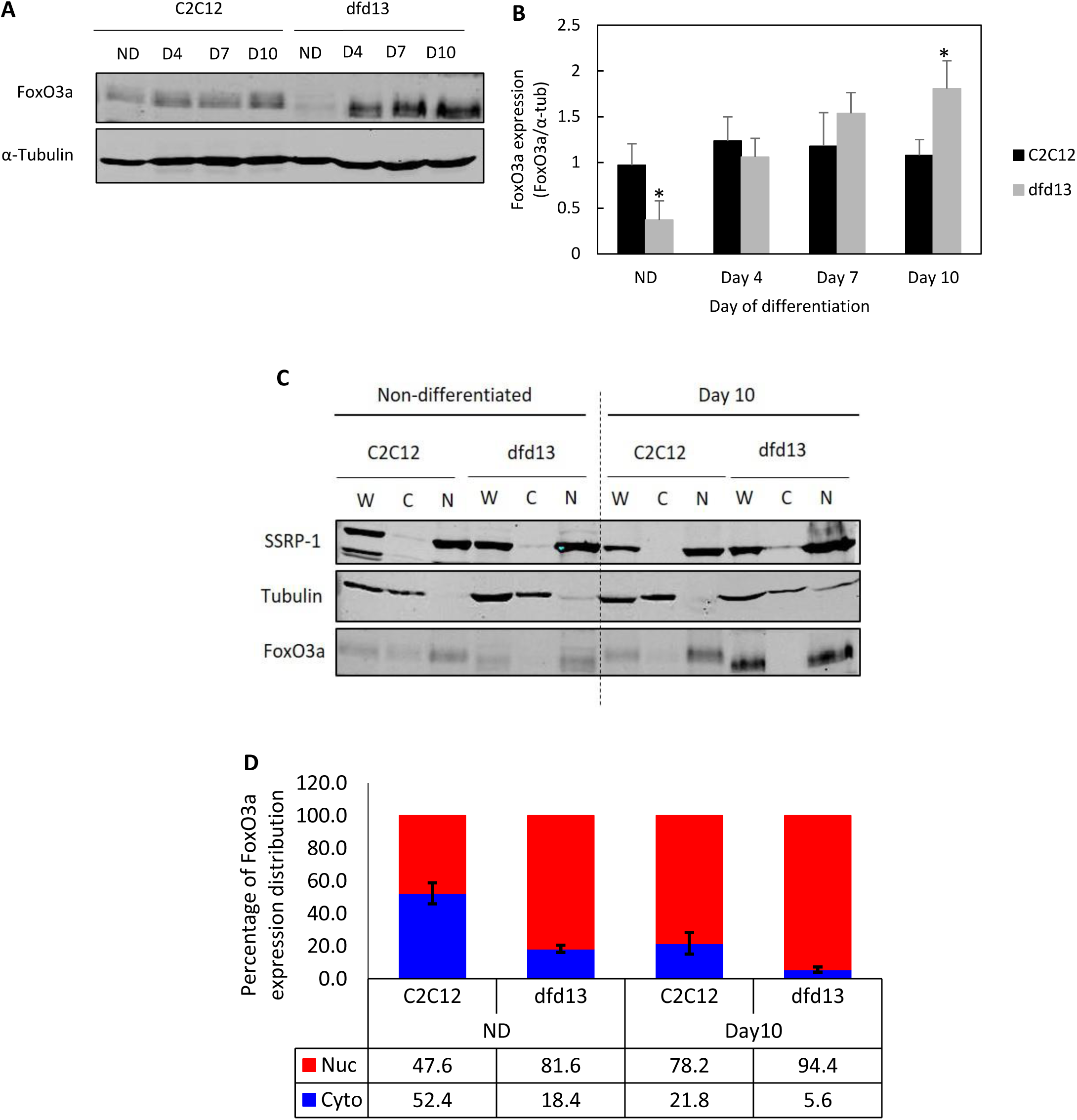
FoxO3a is highly increased and predominantly localised in differentiating dystrophin-deficient myoblasts. Myoblasts were cultured in GM until 80 to 90% confluent before washing twice with PBS and culturing in DM for 10 days with the media being changed every 2 days. Total protein was extracted at the indicated time points prior to immunoblotting with antibodies recognising FoxO3a. (A) Immunoblot analysis of proteins during myoblast differentiation with α-tubulin expression as a loading control. Densitometry analysis representing (B) FoxO3a expression. Subcellular protein extraction was performed at the indicated time points using the REAP protocol prior to immunoblotting with an antibody which recognises FoxO3a. (C) Immunoblot of the proteins during myoblast differentiation with α-tubulin and SSRP1 expression as loading controls. Densitometry analysis representing (D) FoxO3a expression in the cytoplasm and nucleus. The graph represents an average of three repeats from different samples. ND: non-differentiated; *: significantly different (p<0.05) compared to dfd13 myoblasts. GM: growth medium (DMEM + 10% FCS); DM: differentiation medium (DMEM + 2% horse serum). W: whole cell lysate; C: cytoplasm; N: nucleus.

Akt plays a role in inhibiting FoxO3 through phosphorylation at Thr24, Ser256 and Ser319, which leads to nuclear exclusion and activation. As Akt is inactivated in dfd13 myoblasts, it is thought that FoxO3 is not phosphorylated. Therefore, the non-phosphorylated FoxO3 is translocated to the nucleus and activates the gene involved in autophagy. Akt represses FoxO3 via phosphorylation resulting in nuclear exclusion. As Akt is not activated in dfd13 myoblasts, it was speculated that unphosphorylted-FoxO3 translocates into the nucleus and binds to the promoter, thus up-regulating autophagy related genes such as LC3B, Atg5 and Atg7. Therefore, FoxO3 expression and localisation needed to be examined within subcellular fractions, i.e. the nucleus and cytoplasm.

FoxO3 was found to be localised more to the nucleus of dfd13 myoblasts (Fig 5C), with approximately 81.6% of FoxO3 present in the nucleus of dfd13 myoblasts compared to only ∼47.6% in C2C12 myoblasts during the undifferentiated stage. Surprisingly, on day 10 of differentiation, levels in the nucleus had accumulated in both types of myoblast; C2C12 ∼78.2%, and dfd13 ∼94.4% (Fig 5D).

### Autophagy related proteins are highly increased in differentiating dystrophin-deficient myoblasts

It has been shown that FoxO3 is highly expressed and primarily localised to the nucleus of differentiating dfd13 myoblasts. Within the nucleus, FoxO3 acts as a transcriptional activator that can recognise and bind to DNA sequences resulting in the activation of genes involved in autophagy, such as those for Beclin1 and Atgs (30). It was hypothesised that autophagy is highly activated in differentiating dystrophin-deficient myoblasts.

presents immunoblot and densitometry analyses of autophagy related proteins involved in autophagosome formation. Beclin1 expression was slightly increased in C2C12 myoblasts and highly increased in dfd13 myoblasts upon differentiation (Fig 6A). Densitometry analysis of Beclin1 expression showed significant accumulation on day 4 (p<0.05; p = 2.07×10^−2^), day 7 (p<0.05; p = 1.56×10^−2^) and day 10 (p<0.01; p = 5.70×10^−3^) in dfd13 myoblasts compared to C2C12 myoblasts (Fig 6B).

**Fig 6.**
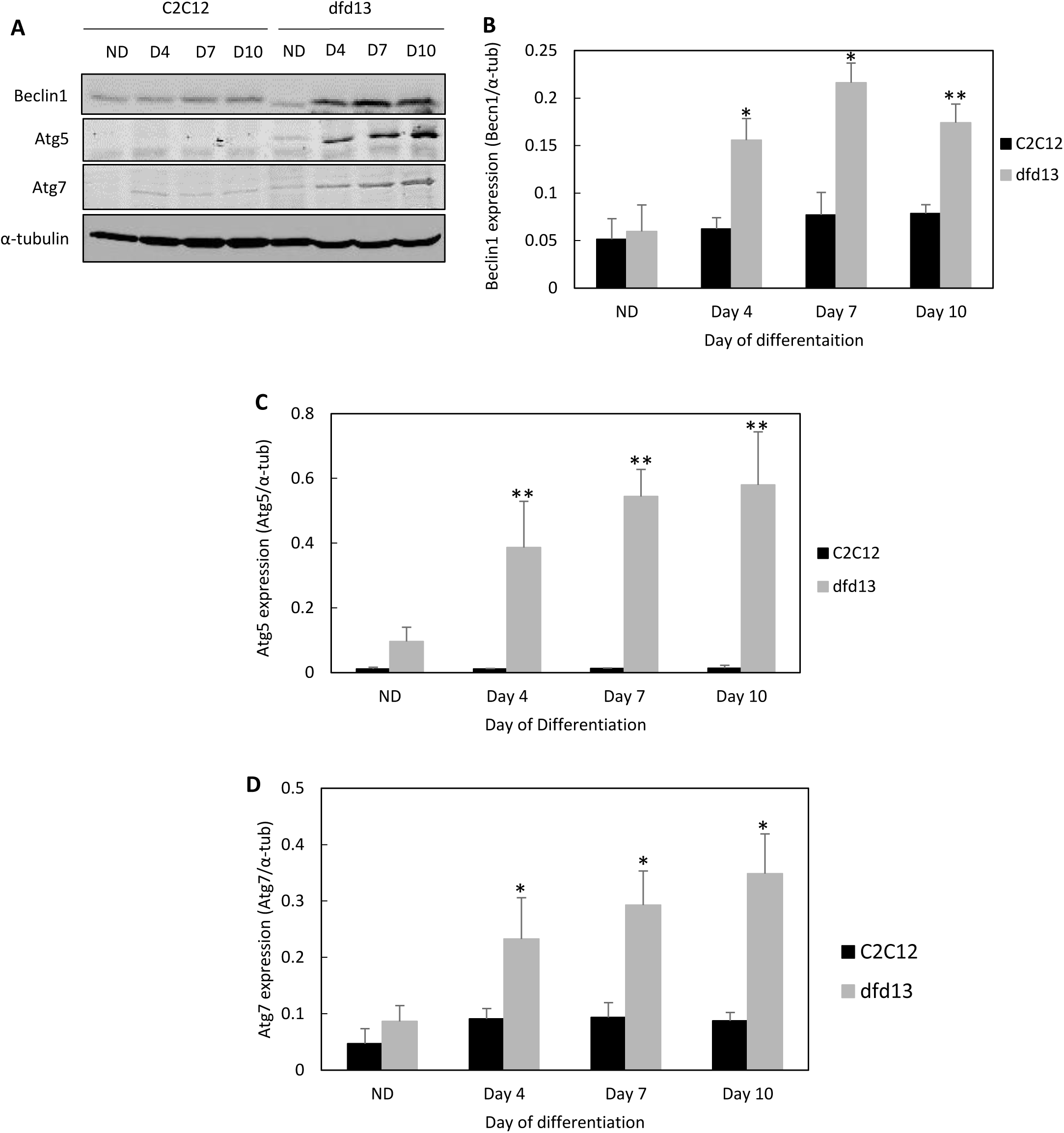
Expression of autophagy related proteins is highly increased in differentiating dystrophin-deficient myoblasts. Myoblasts were cultured in GM until 80 to 90% confluent before washing twice with PBS and culturing in DM for 10 days with the media being changed every 2 days. Protein extraction was performed at the indicated time points. (A) Immunoblot analysis of Beclin1, Atg5 and Atg7 expression with α-tubulin expression as a loading control. Densitometry analysis representing (B) Beclin1, (C) Atg5, and (D) Atg7 expression. The graphs represent an average of four repeats from different samples. ND: non-differentiated; Significantly different: * (p<0.05) and ** (p<0.01) compared to C2C12 myoblasts.

Atg5 was found to be not/less expressed in C2C12 myoblasts (Fig 6C). It can be seen that Atg5 expression was increased upon differentiation in dfd13 myoblasts, and expression was significantly increased on day 4 (p<0.01; p = 6.36×10^−3^), day 7 (p<0.01; p = 6.13×10^−4^) and day 10 (p<0.01; p = 1.96×10^−3^) compared to the non-differentiated stage. Densitometry analysis also showed significant differences (p<0.01) for dfd13 myoblasts at every time point compared to C2C12 myoblasts (Fig 6C).

Densitometry analysis of Atg7 expression in dfd13 myoblasts showed its accumulation throughout the differentiation period; however, there was no significant difference in expression when compared to the non-differentiated stage in C2C12 myoblasts. Expression showed a significant increase in dfd13 myoblasts when compared to the non-differentiated stage and also significant on day 4 (p<0.05; p = 1.43×10^−2^), day 7 (p<0.05; p = 4.54×10^−2^) and day 10 (p<0.05; p = 2.74×10^−2^) compared to C2C12 myoblasts (Fig 6D).

Autophagy is responsible for the removal of unfolded protein as well as dysfunctioning organelles, and has been reported to be constantly active within skeletal muscle (9). Beclin1 is the protein responsible for the initiation of autophagosome formation and is also known as phagophore, while Atg5 and Atg7 are ubiquitin-like enzymes involved in the autophagosome elongation process. In this study, Beclin1, Atg5 and Atg7 showed enhanced expression in dfd13 myoblasts. However, in C2C12 myoblasts there was only a slight increase in the expression of Beclin1 and Atg7 which persisted throughout the differentiation period. Consequently, autophagosome formation (maturation) and autophagic flux during myoblast differentiation were investigated via immunoblotting and flow cytometry analysis, respectively.

### Microtubule-associated light chain-3b expression is increased but autophagic flux is decreased during dystrophin-deficient myoblast differentiation

Autophagy related genes had been shown to be highly activated, and the next step was to determine whether a double-membraned vesicle, known as autophagosome, had been formed. The conversion of Light Chain-3B (LC3B-I) to LC3B-II can be considered to represent total autophagosome formation in myoblasts during differentiation.

As shown in Fig 7A, expression of LC3B-I and LC3B-II is increased in both differentiating C2C12 and dfd13 myoblasts when compared to the respective non-differentiated myoblasts. LC3B-I expression in differentiated dfd13 myoblasts was significantly higher (p<0.01; p = 3.41×10^−3^) at day 10 when compared to differentiated C2C12 myoblasts (Fig 7B). LC3B-II was found to accumulate upon differentiation in both types of myoblasts, but was higher in dfd13 myoblasts, with a significant difference on day 10 (p<0.05; p = 1.69×10^−2^) when compared with to C2C12 myoblasts (Fig 7C). However, the LC3B-II/LC3B-I ratio showed a reduction in dfd13 myoblasts compared to C2C12 myoblasts upon differentiation, and was significantly different (p<0.05; p = 5.33×10^−2^) (Fig 7D).

**Fig 7.**
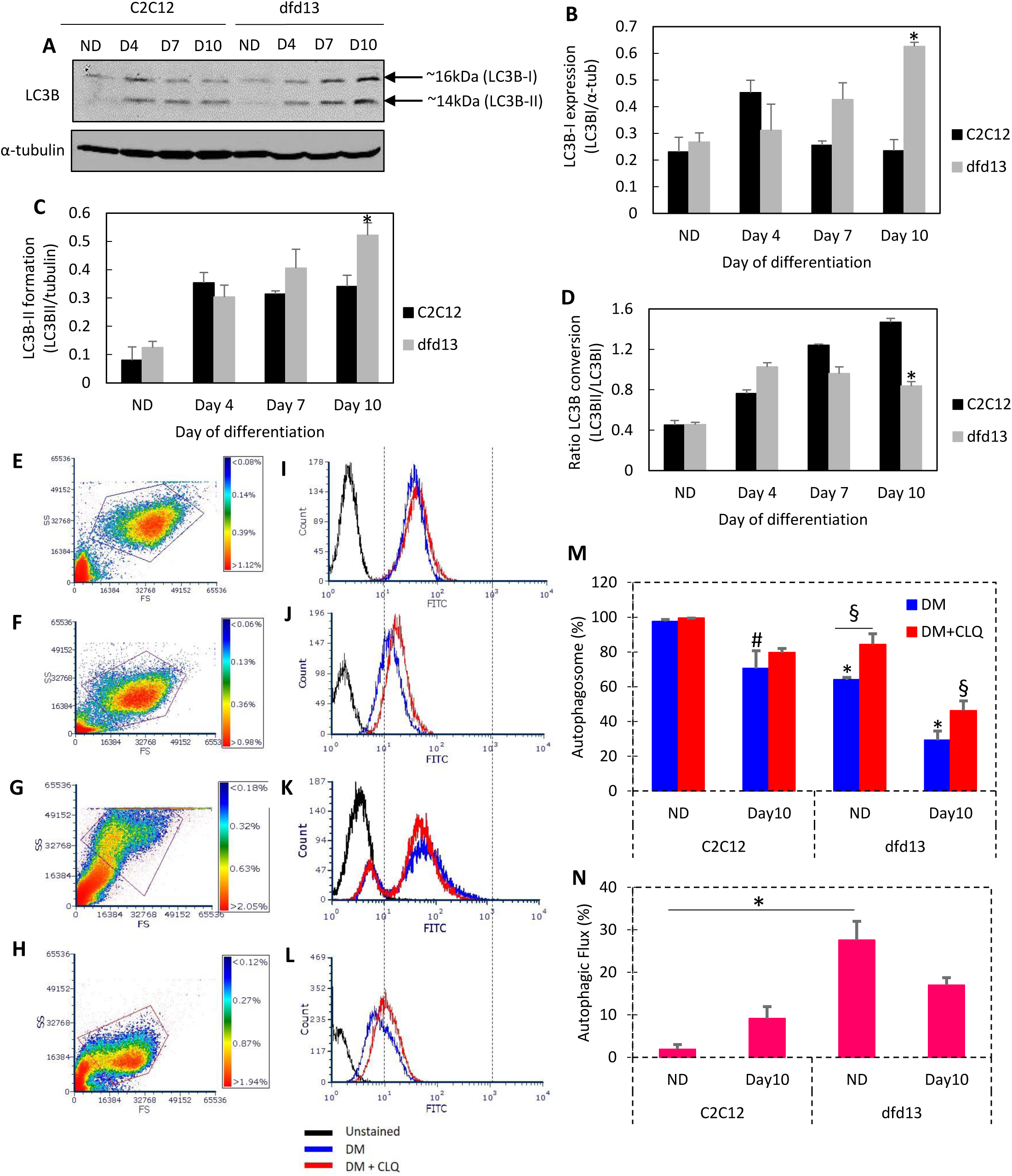
LC3B-II expression is increased but autophagic flux is decreased in differentiating dystrophin-deficient myoblasts. Myoblasts were cultured in GM until 80 to 90% confluent before washing twice with PBS and culturing in DM for 10 days with the media being changed every 2 days. Total protein was extracted at the indicated time points prior to immunoblotting with an antibody recognising LC3B with α-tubulin expression as a loading control. Densitometry analysis of (B) LC3B- I expression, (C) LC3B-II expression, and (D) the ratio of LC3B-II/LC3B-I. For flow cytometry analysis, three different conditions were set up for each stage: 1) unstained cells (negative control); 2) stained cells; and 3) stained cells with chloroquine treatment (positive control). Chloroquine treatment was consisted of 3 hours’ incubation at 37°C. Cells were trypsinised and incubated with Cyto-ID Green stain solution prior to analysis using the FITC channel of a CyAn B flow cytometer (Beckman Coulter, USA). Data analysis was performed using FCS Express 6 Plus Research Edition De Novo Software, USA). The density plot images represent non-differentiated (E) C2C12 myoblasts, (F) dfd13 myoblasts; day10 differentiation (G) C2C12 myoblasts and (H) dfd13 myoblasts. The histogram overlays (I-L) represent each neighbouring dot plot. (M) The percentage of autophagosomes detected in myoblasts and (N) percentage of autophagic flux within myoblasts. All bar graphs represent an average of three repeats from different samples. ND: non-differentiated; significantly different: * (p<0.05) and ** (p<0.01) compared to C2C12 myoblasts. # (p<0.05) when compared with ND; § (p<0.05) when compared to DM+CLQ.

LC3B-II levels correlate with the number of autophagosomes formed; however, this could indicate either the up-regulation of autophagosome formation or a blockage in autophagic degradation. In addition, it does not conclusively indicate autophagic degradation. Therefore, further analysis was performed to determine autophagic flux, with a lysosome inhibitor used as a positive control.

In order to accurately determine autophagic activity, measuring the increase in the number of autophagosomes is required. Previous data showed that LC3B conversion was increased in C2C12 but reduced in dfd13 myoblasts upon differentiation; however, the results from measuring LC3B conversion alone could be inappropriately interpreted. LC3B-II itself has been reported to be degraded by autophagy and also tends to be more sensitive than LC3B-I during immunoblotting analysis (31). Autophagosome formation is an intermediate stage of autophagy, and there could be either the generation of autophagosomes or the blocked conversion of autolysosomes. Therefore, the accurate measurement of autophagic flux is required. In this study a Cyto-ID autophagy detection kit was utilised which selectively labelled autophagic vacuoles independent of the LC3B protein, thus eliminating the need for transfection. This detection kit employs a 488nm-exitable green-emitting fluorescent probe to highlight various vacuolar components of autophagy. Chloroquine was used as a control as it passively diffuses into lysosomes and increases the pH, thus inhibiting lysosome function and blocking fusion with autophagosomes to become an autolysosome.

Fig 7 presents the density plots (E-H) and histograms (I-L) from flow cytometry-based profiling for both C2C12 and dfd13 myoblasts in the non-differentiated and differentiated state. From the density plot images, the size and complexity of each myoblasts can be determined. When the laser beam strikes the stream, a photon passes through unobstructed, but when it strikes a cell there are two different sizes of angle produced. When a photon strikes the cytoplasm, which is less dense and translucent, a small angle of reflection is generated and which can be detected by the forward scatter detector. In contrast, when a photon strikes an organelle such as nucleus or ER (denser), then a wide angle of reflection is generated which is detected by side scatter detector. The forward scatter (FS: X-axis) is proportional to the size, meaning that the bigger a cell the more light that will be scattered and a higher signal will be detected. The side scatter (SS: Y-axis) is proportional to the complexity of the myoblasts.

Based on the density plot images, it can be seen that the dfd13 myoblasts (Fig 7F) are smaller and less complex than C2C12 myoblasts (Fig 7E), although the differences between these cells are not obvious. After differentiation, it can be seen that C2C12 cells become more complex (Fig 7G) while dfd13 cells remain the same (Fig 7H). As both myoblasts were labelled with FITC (green fluorescence), this analysis was taken beyond the basic characteristics of myoblasts, as the specific protein behaviour of the antibody-tagged protein can be accurately measured. The signal was collected and measured/read by using a specific filter/channel based on the size of the emission and excitation of the fluorochrome used.

A Cyto-ID autophagy detection kit and FITC was utilised to label/stain autophagosomes. The collected data was gated and analyses were performed based on the myoblasts in the gated region. Histograms were then plotted, as shown in Fig 7I-J; the black line indicates the unstained myoblasts (negative control), the blue line stained myoblasts, and the red line stained myoblasts treated with chloroquine (positive control). From the histograms data were gated based on the intensity (10^1^-10^3^) (X-axis) and then the percentage obtained.

Generally, both treated and non-treated myoblasts showed a reduction in autophagosome counts after 10 days of differentiation. Non-treated myoblasts had less autophagosomes when compared to treated myoblasts, which is due to chloroquine acting as a lysosome inhibitor, preventing autophagosome from fusing with lysosomes and causing the accumulation of autophagosomes. As illustrated in Fig 7M, the number of autophagosomes was significantly decreased after 10 days of differentiation in both non-treated C2C12 myoblasts (p= 1.98×10^−2^) and dfd13 myoblasts (p= 1.59×10^−2^) when compared to non-differentiated stage. It was also significantly decreased in treated C2C12 myoblasts (p= 2.34×10^−3^) and dfd13 myoblasts (p= 3.44×10^−3^) after 10 days of differentiation. There was a significant reduction in autophagosomes in dfd13 myoblasts when compared to C2C12 myoblasts at the equivalent stages and treatment.

From these results autophagic flux can be determined. Autophagic flux can be calculated by subtracting the chloroquine-treated from the untreated myoblasts, which enables the total number of non-fused autophagosomes to be measured and also the total number of autophagosomes formed. As depicted in Fig 7N, autophagic flux was increased in C2C12 myoblasts after 10 days of differentiation, whereas it was reduced in dfd13 myoblasts. When compared to non-differentiated C2C12 myoblasts, the number of autophagosomes in dfd13 myoblasts was significantly higher (p= 3.64×10^−2^) and was slightly higher in differentiated dfd13 myoblasts compared to differentiated C2C12 myoblasts.

Overall, autophagy activity was decreased upon differentiation in dystrophin-deficient myoblasts. Based on the flow cytometry analysis, autophagic flux was decreased, as the total number of autophagosomes detected was higher in dfd13 myoblasts (both non-differentiated and differentiated) when compared to C2C12 myoblasts. In addition, autophagy was 2-fold higher in non-differentiated myoblasts (proliferation state) compared to the differentiated state.

## Discussion

### Dystrophin-deficient myoblasts do not achieve terminal differentiation

In this experiment it has been established that dystrophin-deficient myoblast differentiation is impaired when in low mitogen medium for 10 days based on morphological analysis (Fig 1A and B) and the detection of F-MyHC expression (Fig 1E). Differentiation analyses showed that less myotubes were formed, and no/less F-MyHC expression was detected. However, pan-myosin was only expressed at the end of differentiation on day 10 in dfd13 myoblasts. Desmin expression levels were found to be higher at the end of the differentiation period. Desmin is a marker for newly formed fibres and is a class-III intermediate filament found in muscle cells that connect myofibrils to each other, as well as to the plasma membrane of myofibres. In this study, it can be suggested that the integrative element in myofibres was generated/developed earlier than myotube formation in the differentiating dystrophin-deficient myoblasts, although full/terminal differentiation was not achieved.

During myogenesis, proliferating myoblasts withdraw from the cell cycle and differentiate into myotubes. Cyclin-dependent Kinase (CDK) inhibitor p21 and retinoblastoma protein (Rb) have been shown to play a critical role in establishing the post-mitotic state by permitting the transcription of the S-phase promoting gene during myogenesis (32). Most mitogens have been shown to promote myoblast proliferation. In contrast, insulin-like growth factors (Igf-1 and Igf-2) trigger myoblast differentiation. It is unknown how Igf-2 is controlled during the initiation of differentiation. A previous study showed that Igf-2 up-regulates its own gene expression via the Akt/PKB pathway (33). Elevation of Igf-2 mRNA was found in C2 myoblasts after 24 hours’ culture in differentiation medium containing 2-10% horse serum (34), while Erbay et al. (2003) suggested that mTOR regulates Igf-2 production at the transcriptional level when cultured under low mitogen conditions (35). There has also been a study suggesting that myoblasts under low mitogen conditions have elevated levels of IGFBP5, which helps concentrate IGF-2 to a threshold level which triggers the IGF-1R pathway leading to differentiation (36,37). However, in this study levels of Igf-1/2 could not be examined in the culture system utilised, i.e. Igf-1/2 secreted/present in the media. Therefore, further study needs to be undertaken to examine this aspect. a previous study by Chen (2013) showed that Igf-2 induction promotes dystrophin-deficient myoblast differentiation for 4 days, and differentiation was greater when induced by a combination of Igf-1/Igf-2/LIF factors (27).

### Elevation of PTEN affects PI3K/AKT regulation in dystrophin-deficient myoblasts

The PI3K/Akt pathway is a highly-conserved pathway for the regulation of skeletal muscle growth and is activated by the binding of Igf-1 to its receptor, Igf1-r. This binding leads to intrinsic tyrosine kinase activation, as well as auto-phosphorylation, which can generate a docking site for IRS. Phosphorylated-IRS then becomes a docking site for PI3K, thus activating PI3K/Akt signalling for myoblast differentiation via a series of phosphorylation events. This signalling is negatively regulated by PTEN.

In this study PTEN was shown to be elevated in dfd13 myoblasts, which had an effect on PI3K activation, as it was found to be decreased during differentiation (Fig 2D), indicating that PI3K regulation is altered in dfd13 myoblasts. PTEN acts as lipid phosphatase and plays a role in removing the phosphate group present on the inositol ring of PIP_3_ to produce PIP_2_. In contrast, PI3K reverses this event by phosphorylating PIP_2_ to become PIP_3_.

PI3K exists as a heterodimer consisting of two subunits; regulatory and catalytic. When cells are stimulated, p85 binds to tyrosine-phosphorylated IRS, and phosphorylated-p85 changes the conformation of p110 and thus mediates the p110 subunit to translocate to the membrane and increases its enzymatic activity such as PIP_2_ phosphorylation. PIP_3_ helps to recruit Akt by binding the plekstrin homolog (PH) domain at the N-terminal of Akt to the cell membrane. PTEN in turn acts as a negative regulator by dephosphorylating PIP_3_ to form PIP_2_, and thereby reduces the available docking sites for Akt to bind to prior to activation. PTEN is mainly found in the cytosol and nucleus. The N-terminal possess a PIP_2_-binding motif while the C-terminal contains a serine/threonine phosphorylation, Ser380, which regulates its stability and activity, as well as membrane recruitment. The main C2 domain, contains basic residues that are essential for membrane binding. Phosphorylation of PTEN is considered result in a closed conformation which is the inactive form. In this conformation, it has been proposed that the phosphorylated C-terminal interacts with the positively charged C2 domain and remains in the cytoplasm (38). As the membrane integrity is disrupted due to the absence of dystrophin, it can be suggested that cytoplasmic-membrane activity is altered in dystrophin-deficient myoblasts and thus there is impaired PTEN-PI3K regulation.

Feron and colleagues (2009) reported that increased PTEN expression caused deregulation of the PI3K/Akt pathway in dystrophin-deficient muscle present in GRMD dogs. This was also observed in muscle sections from 3- to 36-month old animals and indicates that the PI3K/Akt pathway is a long-term alteration. A more recent study by Alexander et al. (2014) proposed that overexpression of micro-RNA-486 improved muscle physiology and performance in dystrophin-deficient mice (39). Micro-RNA-486 is a muscle-enriched micro- RNA that is markedly reduced in the muscle of dystrophin-deficient mice and DMD patient muscles (40).

### Inactivation of Akt is caused by impaired rictor-mTOR2 in dystrophin-deficient myoblasts

In this study, it was found that Akt is not/less activated in dfd13 myoblasts, as phosphorylation at Ser473 or Thr308 was not detected. However, in C2C12 myoblasts Akt was phosphorylated at Ser473 and the activated form accumulated upon differentiation. Immunofluorescence analysis showed that phosphorylated-Akt (Ser473) is localised to the membrane of C2C12 myotubes but found less often in dfd13 myoblasts. During biosynthesis, nascent Akt is phosphorylated at Thr450 within the turn motif site and localised to the cytosol when in its inactive conformation. In the presence of signals via PI3K activation, Akt is recruited to the membrane through the binding of plekstrin homolog (PH) to PIP_3_. Once bound, the conformation is changed and this event unmasks two residues for phosphorylation, Ser473 and Thr308. In this study, less Akt-phosphorylated at Ser473 was detected in dystrophin-deficient myoblasts, indicating that Akt is inactivated/less activated. According to Sarbassov et al. (2005), PDK1 has a better target on Akt phosphorylated at Ser473 than non-phosphorylated Akt (25). In line with the results shown Fig 3, there is no Thr308 phosphorylation if Ser473 phosphorylation does not occur. However, the expression of total Akt did not show any significant difference when compared between both types of myoblast.

Since Akt was found to only be phosphorylated at Ser473 the protein responsible for this phosphorylation event was investigated. It is known that mTORC2, specifically, rictor, can directly phosphorylate Akt at Ser473 *in vitro* and this facilitated Thr308 phosphorylation by PDK1 in drosophila (25). In this study, it was found that rictor is inactivated in both undifferentiated and differentiated dfd13 cells, which explains the absence of phosphorylated-Akt (Ser473) observed; however, the regulator for rictor activation in myoblasts remains unknown. Rictor phosphorylation has been shown to require mTORC1-activated p70S6K in MEF, HEK293 and HeLa cells (41). Recently, it has been showed that PTEN negatively regulates mTORC2 signalling in glioma (42). As PTEN was found to be highly expressed in dfd13 myoblasts, inactivation of rictor can be seen as a consequence of this event and presumably represents the same scenario as in myoblasts.

### Excessive formation of autophagosome in dystrophin-deficient myoblast via FoxO3-mediated regulation

In skeletal muscle autophagy is transiently activated and continues for only a few days (9,43); it is regulated via FoxO3. Generally, it can be seen that autophagosomes are formed in both types of myoblast during differentiation, as the autophagy related protein involved in the initiation and elongation process was found to be increased. The ratio of LC3B-I converted to LC3B-II also reflected the formation of autophagosomes during differentiation (Fig 7D).

In this study, it can clearly be seen that autophagosome formation is modulated via FoxO3 in dfd13 myoblasts. Inactivation of Akt allowed the active-form of FoxO3 (unphosphorylated) to translocate into the nucleus and trigger the expression of autophagy related genes. FoxO3 is required for the transcriptional regulation of LC3B. and also for transcriptional regulation of MAFbx and MuRF1. This transcriptional regulation leads to protein degradation via the autophagy-lysosome pathway and ubiquitin-proteosome pathway, respectively. However, autophagy in C2C12 myoblasts seems to be only partially FoxO3- mediated, as subsequent activation of Akt inhibited FoxO3 and suppressed its translocation to the nucleus and the targeting of autophagy-related gene activation. Activated-Akt, through phosphorylation at Ser473 by rictor-mTORC2, also contributes to autophagy activation during C2C12 myoblast differentiation (28,44).

### Modulation of Atg5-dependent autophagy during differentiation of dystrophin-deficient myoblasts

Autophagy is responsible for removing unfolded proteins as well as dysfunctioning organelles, and has been reported to be constantly active within skeletal muscle. Recently, increased autophagy has been reported to protect differentiating myoblasts from apoptotic cell death (45). Several autophagy related genes are known to be involved in the formation of autophagosomes, and FoxO3 has been shown to induce multiple autophagy related genes, including LC3B transcription in skeletal muscle (30). LC3B is an isoform of LC3 which plays a critical role in autophagy via post-translational modification.

LC3B is a subunit of microtubule-associated protein 1 (MAP1LC3B), and LC3 is cleaved by Atg4 to become cytosolic LC3B-I. LC3B-I is then converted to lipidated-LC3B-II through the conjugation of membrane lipid phosphatidylethanoleamine (PE), which involves the E1-like enzyme ubiquitin, Atg7 and the E2-like enzyme, Atg10. LC3B-II then binds to the isolation membrane and mediates membrane elongation until the edges fuse to form an autophagosome. The isolation membrane appears when cells are placed under starvation conditions. Atg7 also conjugate Atg5 to Atg12 to form the Atg5-Atg12 complex and then bind to the isolation membrane with Atg16. This complex binding is necessary for autophagosome formation.

In this study, the results showed that LC3B-I was increased in both types of myoblast due to the up-regulation of Beclin1 by nuclear-FoxO3, which is increased in dystrophin-deficient myoblasts. Beclin1 forms a complex with Vsp34 and becomes a core component during the pre-autophagosome stage. This complex then binds to the ULK1/Atg13/FIP200/Atg101 complex before entering the elongation stage, when Atg7 catalyses the ligation of Atg5 to Atg12 to become the Atg5-Atg12 complex. With the aid of the Atg5-Atg12 complex, Atg7 catalyses the transfer of PE to LC3B-I, converting it to LC3B-II resulting in the accumulation of LC3B-II within dfd13 myoblasts (Fig 7C). Therefore, autophagosomes can be formed even when Atg5 is not present and so it can be suggested that Atg5 plays a booster role for autophagosome formation in dfd13 myoblasts. Autophagy in C2C12 myoblasts might also activated via PERK-mediated CHOP (46). Although the expression of both LC3B-I and LC3B-II in differentiating dfd13 myoblasts accumulates until day 10, the conversion ratio of LC3B-I to LC3B-II was reduced in dystrophin-deficient myoblasts, indicating that autophagy regulation is impaired.

It has been reported that autophagy is induced upon myoblast differentiation in order to eliminate pre-existing structures and proteins. This elimination occurs concomitantly with the myoblast fusion process prior to the formation of multinucleated myotubes. Based on the results presented, autophagy is activated in both types of myoblasts; however, over expression of Atg7 and Atg5 in dystrophin-deficient myoblasts suggests that the ubiquitin-like system is impaired. This could affect cascade activation which in turn will affect autophagosome formation, as well as autophagic flux in dystrophin-deficient myoblasts.

### Reduction in Autophagic Flux Proves There is Defective Autophagy in Dystrophin-Deficient Myoblasts

From the data obtained the increased expression of autophagy related genes (Atg5, Atg7 & Beclin1) demonstrates excessive autophagosome formation in dfd13 myoblasts during differentiation; however, the conversion ratio of LC3B-I to LC3B-II was reduced in dystrophin-deficient myoblasts. Furthermore, although activation was higher, autophagic flux analysis showed a reduction upon differentiation and revealed that autophagy activity is decreased upon differentiation in dfd13 myoblasts. Therefore, it was suggested that after excessive autophagosome formation in non-differentiated dfd13 myoblasts could potentially undergo apoptosis during differentiation. Previously, our group has shown that the ratio of cleaved caspase-3 to uncleaved caspase-3 is increased in a derivative of dfd13 myoblasts, PD50A when cultured in DM (27). This finding fits with the current data obtained and in the next chapter it is shown that dystrophin-deficient myoblasts are prone to apoptosis via PERK-mediated CHOP under ER stress conditions.

DMD is mostly characterised by ∼2 years of age and it progresses until the early 20s. Therefore, the applicability of this finding to the understating of DMD patients may fit with the actual scenario. During the early stage of life (new born to below 2 years), deficient myoblasts survive as most cells are actively proliferating and high levels of autophagy activation is able to prevent apoptosis. As age increases (>2 years old), most of the deficient myoblasts have been triggered for differentiation as the body develops and muscle size and functionality increases, but now autophagy starts to decrease and apoptosis begins. This is the most common period for when DMD is diagnosed. At the later stage (∼10 years-old), there is progressive disruption to the muscle due to loss/damage and support (wheelchair) is needed for mobility and undertaking daily life tasks. This state will worsen and patients will eventually die, commonly as a complication of respiratory muscle damage and cardiomyopathy.

## Conclusion

PTEN-PI3K/Akt and its downstream proteins are perturbed in dystrophin-deficient myoblasts. From the data obtained, dystrophin-deficient myoblasts exhibit the high expression of autophagy related proteins. However, a reduction in autophagy activity, as well as autophagy flux upon differentiation, indicates that autophagy is defective. This finding suggests a new mechanism for the reduction of autophagy in dystrophin-deficient myoblasts. The perturbation of the PTEN-PI3K/Akt pathway initially triggers excessive autophagosome formation, and subsequently there is a reduction in autophagic flux within dystrophin-deficient myoblasts.

## Acknowledgements

The authors would like to thank to Dr. Chen Hung-Chih (Acedemia Sinica, Taiwan) for editorial and laboratory assistance, Dr. Melissa Grant (School of Dentistry, University of Birmingham), Dr. Zubair Ahmed (Medical School, University of Birmingham), Shabana Begum and Adil Rashid (University of Birmingham) for reagents supplement, Dr. Matthew MacKenzie (Institute for Biomedical Sciences (IBR), University of Birmingham) for flow cytometry data acquisition, Professor John K. Heath (University of Birmingham) and Professor Theodore Fotsis (Institute of Molecular Biology & Biotechnology, University of Ioannina, Greece) for helpful comments and idea throughout the work.

## Authors Contribution

Design the experiments: MDY JS. Performed the experiments: MDY. Analyzed the data: MDY JS. Contribute reagents/materials/analysis tools: JS. Wrote the paper: MDY JS

